# Local and landscape drivers of aquatic-to-terrestrial subsidies in riparian ecosystems: a worldwide meta-analysis

**DOI:** 10.1101/446815

**Authors:** D. Lafage, E. Bergman, R. L. Eckstein, M. Österling, J.P. Sadler, JJ Piccolo

**Affiliations:** Karlstad University. Department of Environmental and Life Sciences / Biology; School of Geography, Earth and Environmental Sciences, The University of Birmingham, Birmingham, United Kingdom

**Keywords:** anthropogenic land use, aquatic subsidies, diet, human population, stable isotopes, terrestrial predators

## Abstract

Cross-boundary fluxes of organisms and matter, termed “subsidies”, are now recognized to be reciprocal and of roughly equal importance for both aquatic and terrestrial systems, even if terrestrial input to aquatic ecosystems has received most attention. The magnitude of aquatic to terrestrial subsidies is well documented, but the drivers behind these subsidies and their utilization by terrestrial consumers are characteristically local scale studies, limiting the inferences that can be drawn for broader geographic scales. We therefore built and analyzed a database of stable isotope data extracted from 21 studies worldwide, to identify both landscape and local scale variables that may affect the diet of terrestrial predators in riparian ecosystems. Our meta-analysis revealed a greater magnitude of aquatic-to-terrestrial subsidies (> 50%) than previously reported, albeit with large geographic and inter-annual variations. We demonstrated a large effect of landscape-scale factors on aquatic-to-terrestrial subsidies, particularly anthropogenic land use and tree cover. Local human population was the only relevant factor at the local scale. We also found that studies on landscape-scale and anthropogenic land use effects on aquatic-to-terrestrial subsidies are currently strongly under-represented in the ecological literature. Such studies are needed to improve our understanding of how land use and environmental change might influence future patterns of biodiversity and ecosystem function.

## Introduction

Decades of research have demonstrated and quantified the tight linkages between aquatic and terrestrial ecosystems (Fisher and Likens 1973, Bartels et al. 2012). Cross-boundary fluxes connecting ecosystems, usually termed “subsidies” (Polis et al. 1997b), can be organisms, energy, or nutrients. Terrestrial-to-aquatic subsidies in the form of litter and organic matter are essential for aquatic ecosystem function (reviwed by Tank et al. 2010) and terrestrial prey subsidies also have important effects on riverine food-webs (Polis and Hurd 1996, Nakano and Murakami 2001, Erős et al. 2012, Gustafsson et al. 2014). More recently, research has focused on reciprocal subsidies between aquatic and terrestrial ecosystems (Baxter et al. 2005, Schindler and Smits 2017). Although the amount of terrestrial-to-aquatic prey subsidies often is greater than the reverse, their overall contribution to the carbon budget of predators is similar (Bartels et al. 2012). Thus, the most recent picture to emerge is that of tightly-coupled, roughly reciprocal aquatic-terrestrial ecosystems, at least at the local scale at which most studies have taken place.

One of the remaining key challenges for understanding the ecology of cross-boundary fluxes is to determine at which scales and to what extent the structure of the surrounding terrestrial landscape affects the magnitude and the importance of aquatic-to-terrestrial subsidies (Marcarelli et al. 2011). At the local scale (100 m buffer), landscape structure has an impact on predator diet by facilitating or preventing subsidies from entering recipient ecosystems (Greenwood 2014, Muehlbauer et al. 2014). At the landscape (catchment) scale, ecosystem size (McHugh et al. 2010, Jackson and Sullivan 2017) and land use (Stenroth et al. 2015, Carlson et al. 2016) have recently more attention. Studies focusing on the effect of ecosystem size and land use on riparian ecosystem food webs, however, remain scarce (e.g. Marczak et al., 2007; Schindler and Smits, 2017). Land use, at local and landscape scales, influences the composition and biomass of both aquatic insect communities (via water quality, terrestrial subsidies and canopy cover: Dolédec et al., 2006; Schindler and Smits, 2017; and predator communities: Hendrickx et al., 2007; Lafage et al., 2015). On the other hand, ecosystem size, by integrating the effects of spatial heterogeneity, disturbance and productivity, is a strong predictor of food chain length (Sabo et al. 2010). To gain a better understanding of broader-scale ecological processes, comparative studies of aquatic-terrestrial ecosystems at the catchment scale are needed.

In this study, we conducted a worldwide meta-analysis of studies that have assessed aquatic-to-terrestrial subsidies using stable isotopes. We quantified the effects of ecosystem size, stream morphology and land use on aquatic subsidies to terrestrial predators. First, we estimated the overall proportion of aquatic subsidies in the diet of several groups of terrestrial predators, and tested whether the proportion of these prey was significantly higher than that of terrestrial prey. We hypothesised that the proportion of aquatic subsidies varied between taxonomic groups of predators, hydrological system type (hydro-ecoregion) and year. Next, we assessed the relative importance of biotic and abiotic variables at local- and landscape- scales (100 m buffers and catchments, respectively) for the proportion of aquatic subsidies in the diet of spider and carabid beetle predators. We hypothesised that landscape-scale variables related to anthropogenic land use would be of at least equal importance in explaining predators’ diets as commonly-assessed local-scale variables.

## Methods

Our meta-analysis focused on the use of aquatic subsidies by terrestrial predators. We restricted the subsidies to aquatic organisms actively crossing the boundary between aquatic and terrestrial ecosystems (i.e. macro-invertebrates). All predators consuming aquatic macro-invertebrates were included. In order to get a more accurate estimation of the proportion of aquatic subsidies in the diet of predators, we restricted our meta-analysis to studies using stable isotopes, which integrate the use of prey types over a longer period of time than do stomach content analyses (Tieszen et al. 1983).

### Data retrieval

We searched the Web of Science and Google Scholar for studies focusing on riparian habitats and using stable isotopes as a tool to infer the contribution of aquatic prey to the diet of terrestrial predators. The keywords used were “aquatic subsidies” AND “stable isotope” AND “diet”, which gave 69 results. From these 69 articles we refined the selection in several steps. First, a selection was made based on words in the title and a second one on words in the abstract. We then screened the bibliography of the selected studies to find new references and iterated this search procedure until we did not find any new documents. This procedure reduced the 69 papers to 47. At last, a selection of studies was based on the number of sampling sites and replicates in the different studies, i.e. we kept studies with at least two sampling sites or studies with repeated measurements in time and studies including sampling of two predator species.

As studies using experimental manipulation of subsidies (and using stable isotopes) were very rare, descriptive studies were also included. Studies on predators’ diet based on stable isotopes include a great variety of techniques used to partition the diet between aquatic and terrestrial prey (mainly linear mixing models *vs* Bayesian mixing models), and great differences in the assumed isotope fractionation between trophic levels. To overcome this issue we (re)-calculated the percentage of aquatic prey in the diet of predators using the same Bayesian mixing model and fractionation values. Using the same fractionation values for all studies was essential as Bayesian mixing models may be highly sensitive to the value used (Bond and Diamond 2011). Consequently, we rejected studies in which the mean and standard deviation of δ13C and δ15N for consumers and prey per sampling site could not be extracted. The final data set consisted of 21 studies (Table 1). Data were retrieved from tables, supplementary material, figures (using WebPlotDigitizer) or by contacting the authors.

**Table 1.**
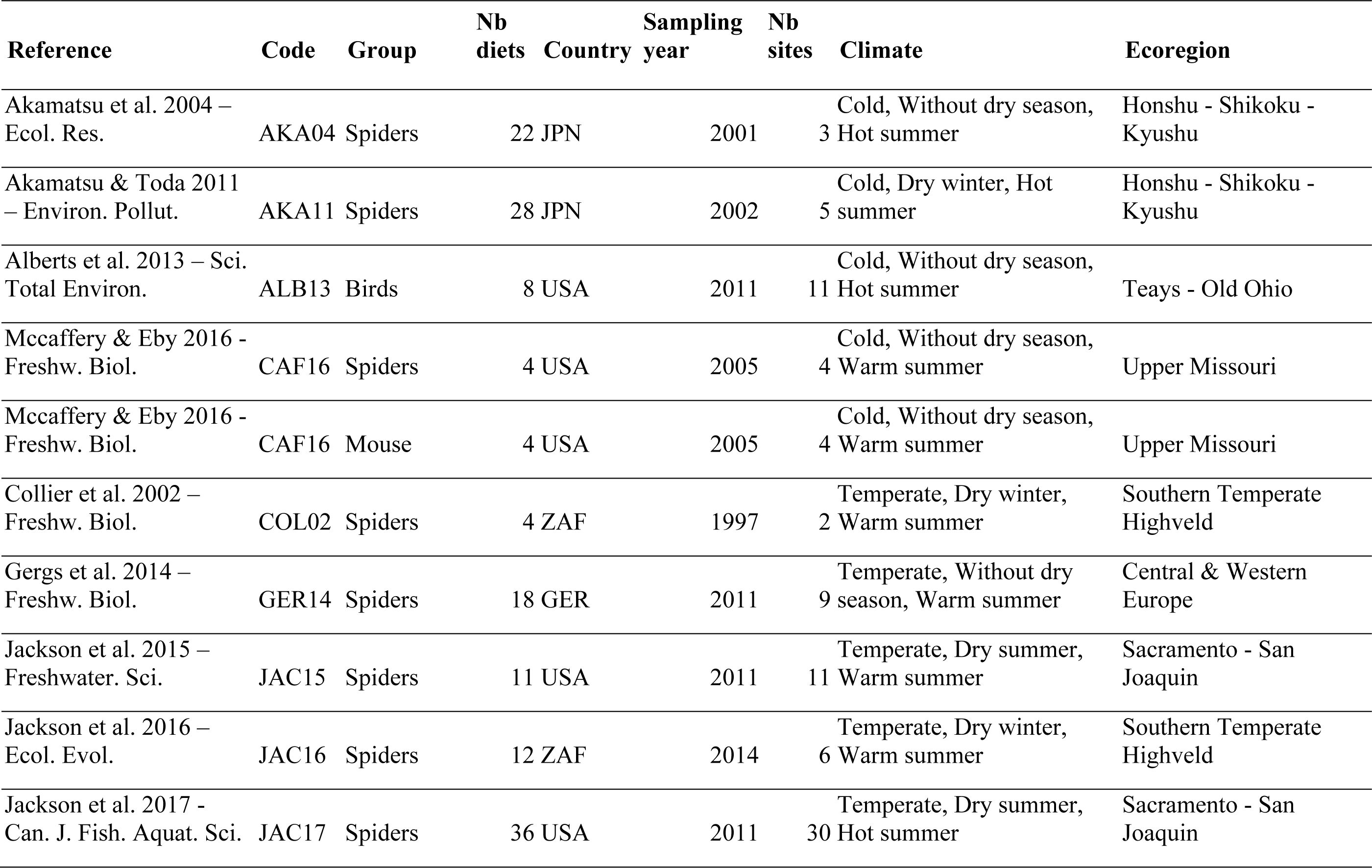
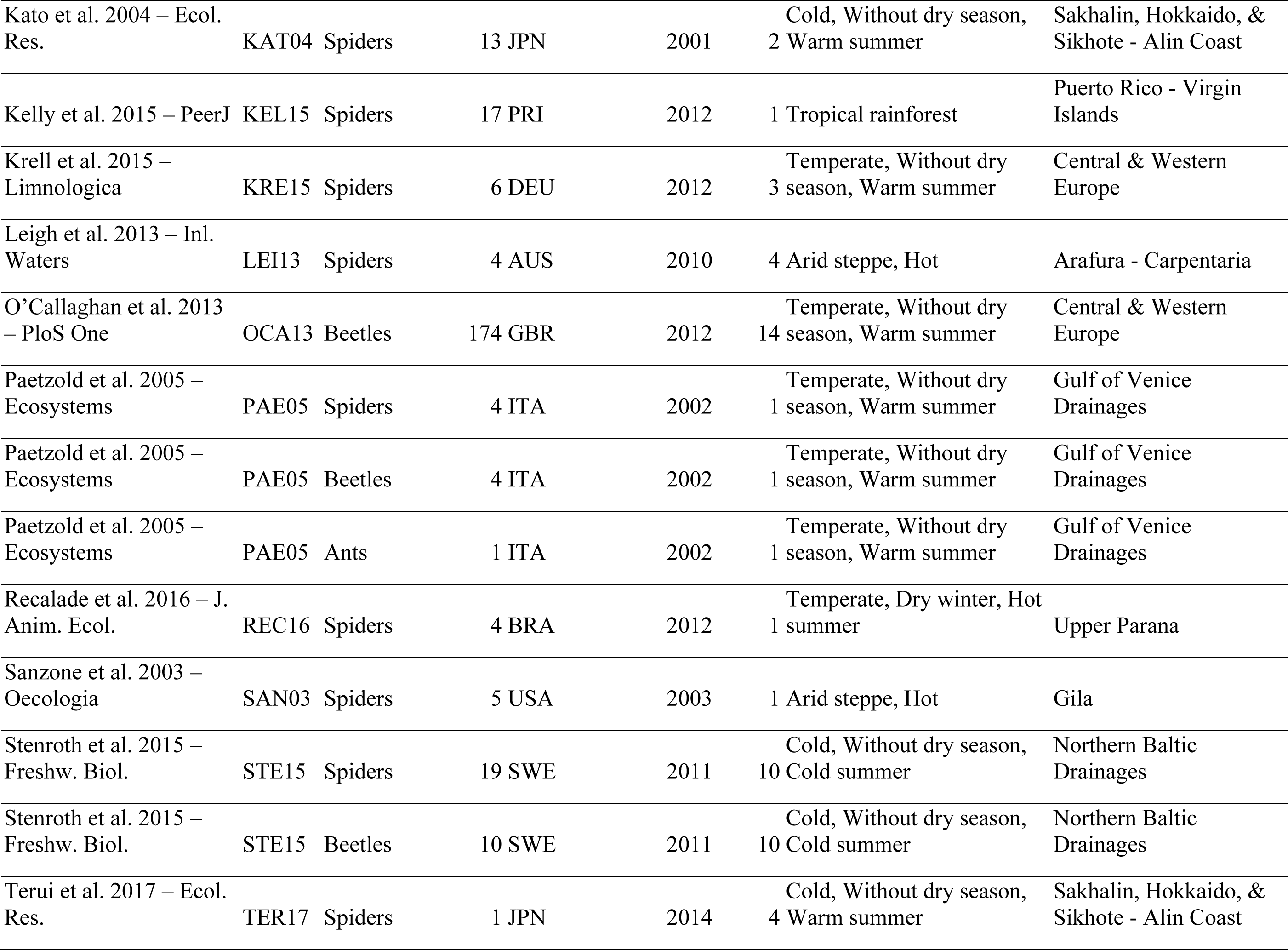
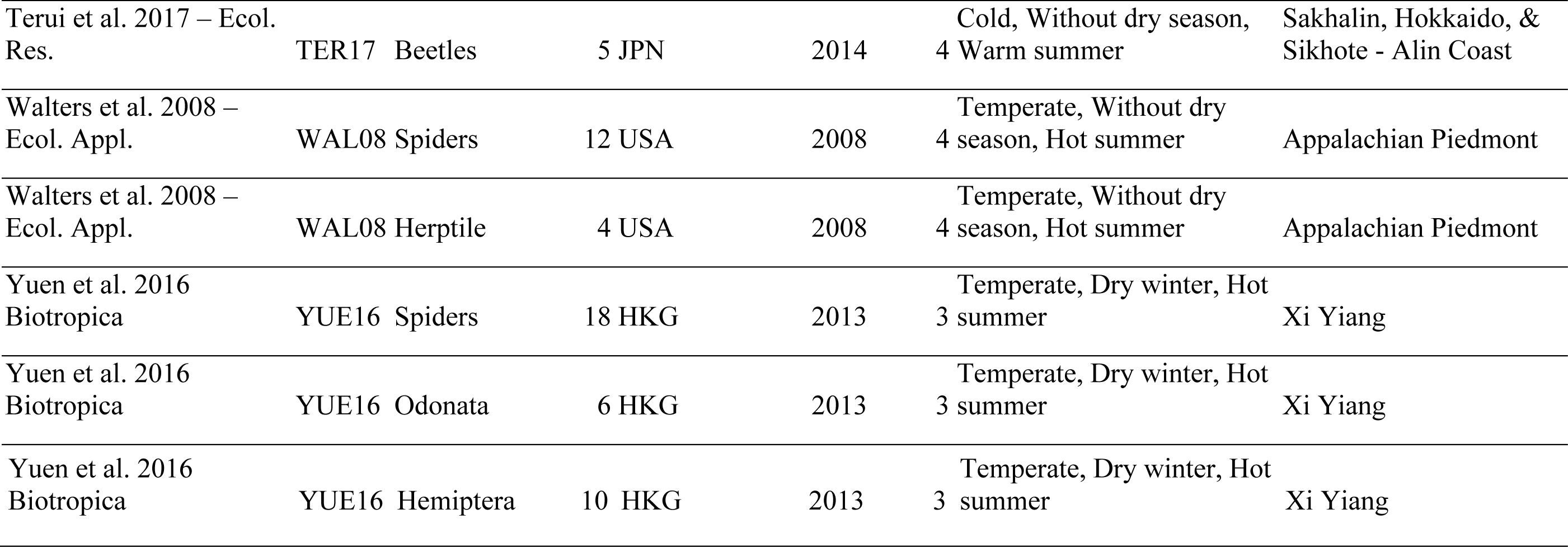
Characteristics of the studies used for the meta-analysis. Climate is extracted from Peel et al. (2007) and ecoregion from Abell et al. (2008)

### Response variable

The proportion of aquatic subsidies in predators’ diet was inferred using two-source Bayesian mixing models. Inputs to the models were means and standard deviations for δ13C and δ15N of aquatic and terrestrial preys with fractionation values recommended by McCutchan et al. (2003). In some studies, δ13C and δ15N values were only available for basal sources (algae and terrestrial litter). In these cases, trophic fractionation was estimated using the per trophic step fractionation multiplied by the estimated number of trophic transfers between the consumer and basal resources. This number was estimated as the difference between the consumer δ15N and mean basal resource δ15N divided by 3.4%o (McHugh et al. 2010, Jackson and Sullivan 2017). When raw data for stable isotope were available for consumers, we used the simmr package (Parnell et al. 2013, Parnell 2016) to infer the proportion of aquatic vs terrestrial subsidies in diet. When only means and standard errors were available we used a modified version of the JAGS models used by Parnell et al. (2013) to include standard error of the consumer isotope values as a prior of the model. Source aggregation (terrestrial vs aquatic) was made *a priori* as the number of sources included in models was variable between studies, which is problematic for *a posteriori* aggregations if one wants to compare diets (Stock et al. 2018). We chose not to give any prior to the proportion of aquatic preys in diet (generalist diets) which means that all possible combinations of proportions of aquatic and terrestrial preys were likely *a priori* (Stock et al. 2018).

### Predictors

The catchment draining to each sampling location was delineated using QGIS 2.18.18 (Quantum GIS Development Team 2017) and GRASS (GRASS Development Team 2017) plugin r.watershed from a 30 m resolution digital elevation model (Shuttle Radar Topography Mission (SRTM) 1 Arc-Second Global, LP DAAC). Predictors were extracted at local (100 m buffer) and landscape (catchment) scales. At the landscape scale, the predictors were catchment perimeter-to-area (a function of size, shape, and fractal irregularity or folding of the edge: Polis et al., 1997a); percentage cover of agriculture, forests, non-forested natural habitats (bare ground, herbaceous, shrubs), open waters (lakes and meadows) and urban areas; mean percent tree cover (a measure of canopy cover); and mean human population. At the local scale, the predictors were river width; meandering ratio over 1 km upstream; land use class; mean percent tree cover; and mean human population.

Land use data were extracted from GLCNMO v3 (Tateishi et al. 2014). Percent tree cover was extracted from PTC V2 (Geospatial Information Authority of Japan, Chiba University and collaborating organizations). Mean human population was extracted from Gridded Population of the World, Version 4 (Center for International Earth Science Information Network, 2016). River width and meandering ratio were extracted under GIS using google maps satellite imagery. To take into account the possible influence of climate, location and local biodiversity, each sampling site was assigned to a freshwater ecoregion according (Abell et al. 2008).

### Statistical analysis

We used the proportion of aquatic subsidies in the diet minus 0.5 as an effect-size to test for differences between proportion of aquatic and terrestrial subsidies in the diet of the terrestrial predators. Freshwater ecoregion, sampling year and taxonomic group of the predators were included in the model as fixed factors. We used the metafor package (Viechtbauer 2010) with restricted maximum-likelihood estimator to test the effect-size.

The selection of landscape and local variables best explaining the proportion of aquatic subsidies in predators’ diet was done using partial least square regression (PLS) on mean % of aquatic subsidies in the diet per sampling site. Given the low number of studies available for some groups (Table 1), the PLS were only performed for spiders and carabid beetles. Freshwater ecoregion and sampling year were also included in the model as moderators. PLS regression extracts orthogonal components (latent variables maximizing the explained variance in the dependent variables) from a set of variables (Eriksson et al. 2006) and are particularly useful when dealing with correlated predictors (Carrascal et al. 2009), which is often the case for land use variables. The number of components to be kept was determined based on Q^2^ value with a M-fold cross-validation approach. Eriksson et al. (2006) recommend a ‘variable importance on the projection’ (VIP) greater than 1 for identifying the most important predictors. Predictors with 0.8<VIP<1 explain only some variation in the model and predictors with 0.8<VIP are considered non-explicative. Weights of the variables (loading values) describe the direction and strength of the relationship between predictor and dependent variables. The PLS were performed using mixOmics package for R (Le Cao et al. 2017). As we expected different scale effects according to taxonomic group, the PLS were performed separately for each group.

Dataset and code are available on the Open Science Framework repository (DOI 10.17605/0SF.I0/T6EYP).

## Results

### Dataset description

The final dataset resulted in 21 studies representing 159 sampling sites and 400 diets (Table 1). This corresponds to almost half of the studies initially selected. Twenty-six studies could not be used, mainly because they did not report data in a suitable format and quality for analysis of diet partitioning. Among these 21 studies, two were not used in the PLS because we could not locate the sampling sites with enough accuracy. Spiders and carabid beetles were the two most studied groups whose diets were estimated in 51.3% and 41.6% of the studies, respectively. The studies were mainly located in the northern hemisphere with cold or temperate climates (Fig. 1 and Table 1).

Study site locations were strongly biased toward small forested catchments with very low human population density and urbanization extent and located mainly in the northern hemisphere (Fig. 2 and 3). Conversely, a few studies were also located in rivers with very large catchments or/and high human population.

**Figure 1:**
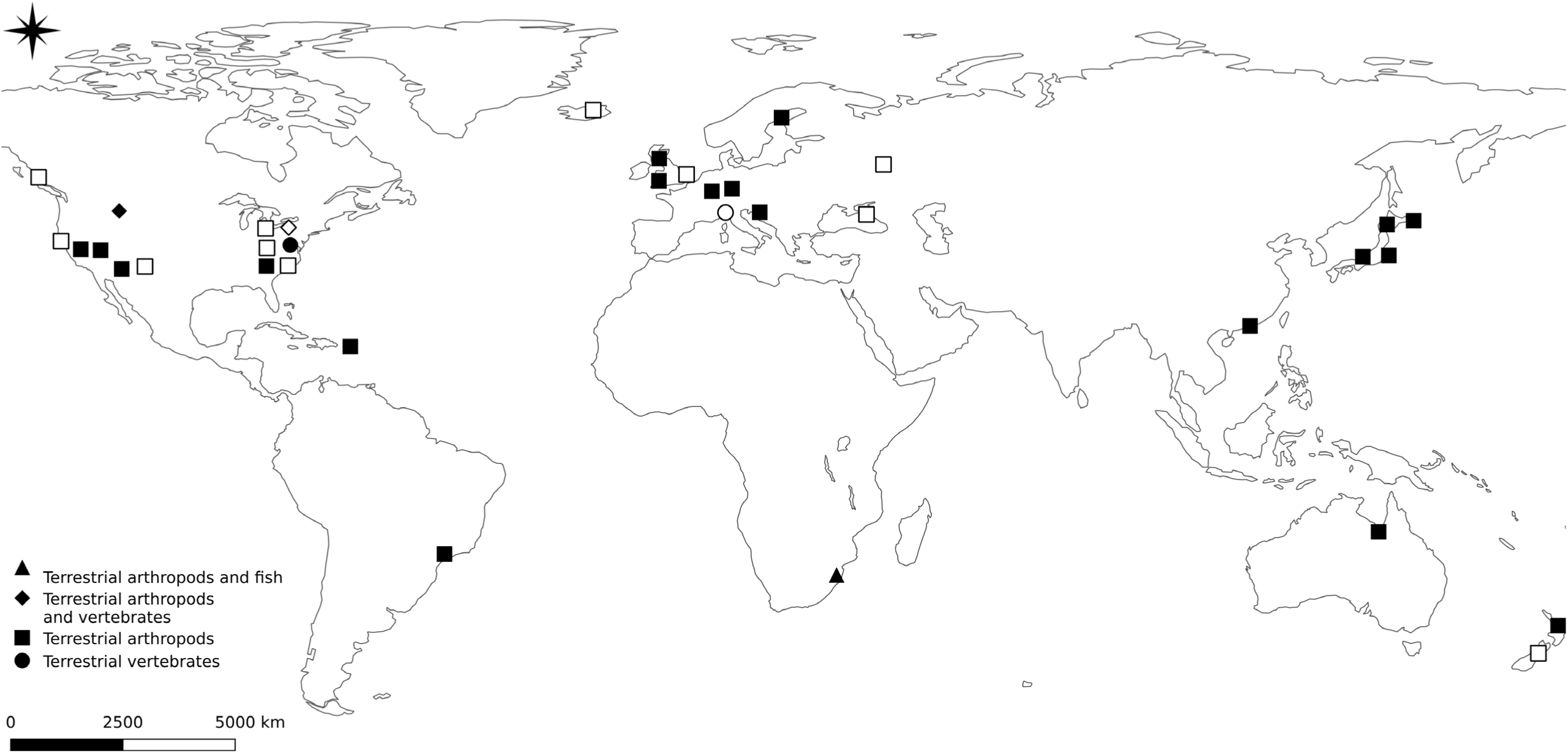
Map of the selected studies. White symbols are studies that were rejected on data quality grounds (see text for details).

**Figure 2:**
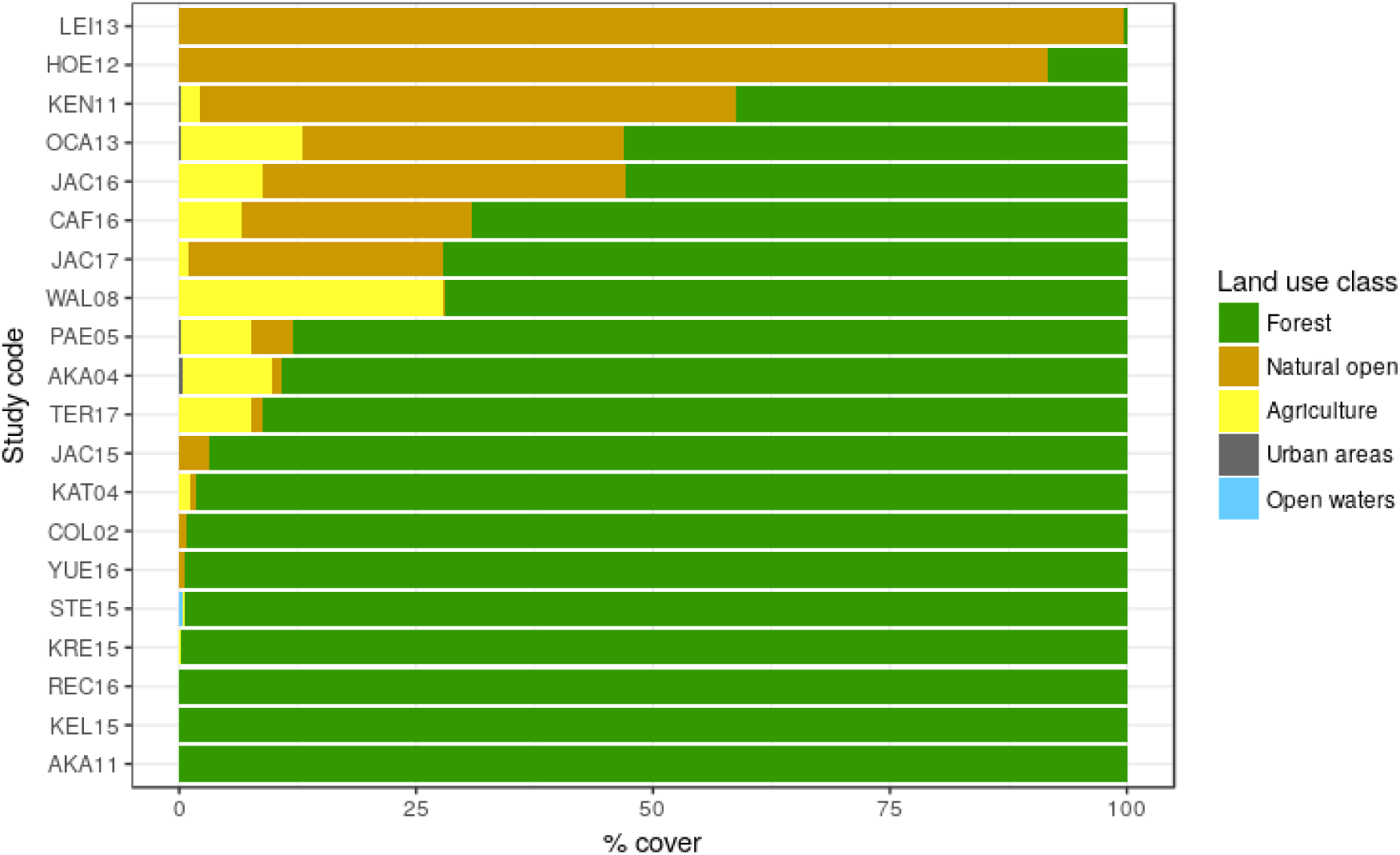
Plot of the percentage cover of each land use class in catchments per study.

**Figure 3:**
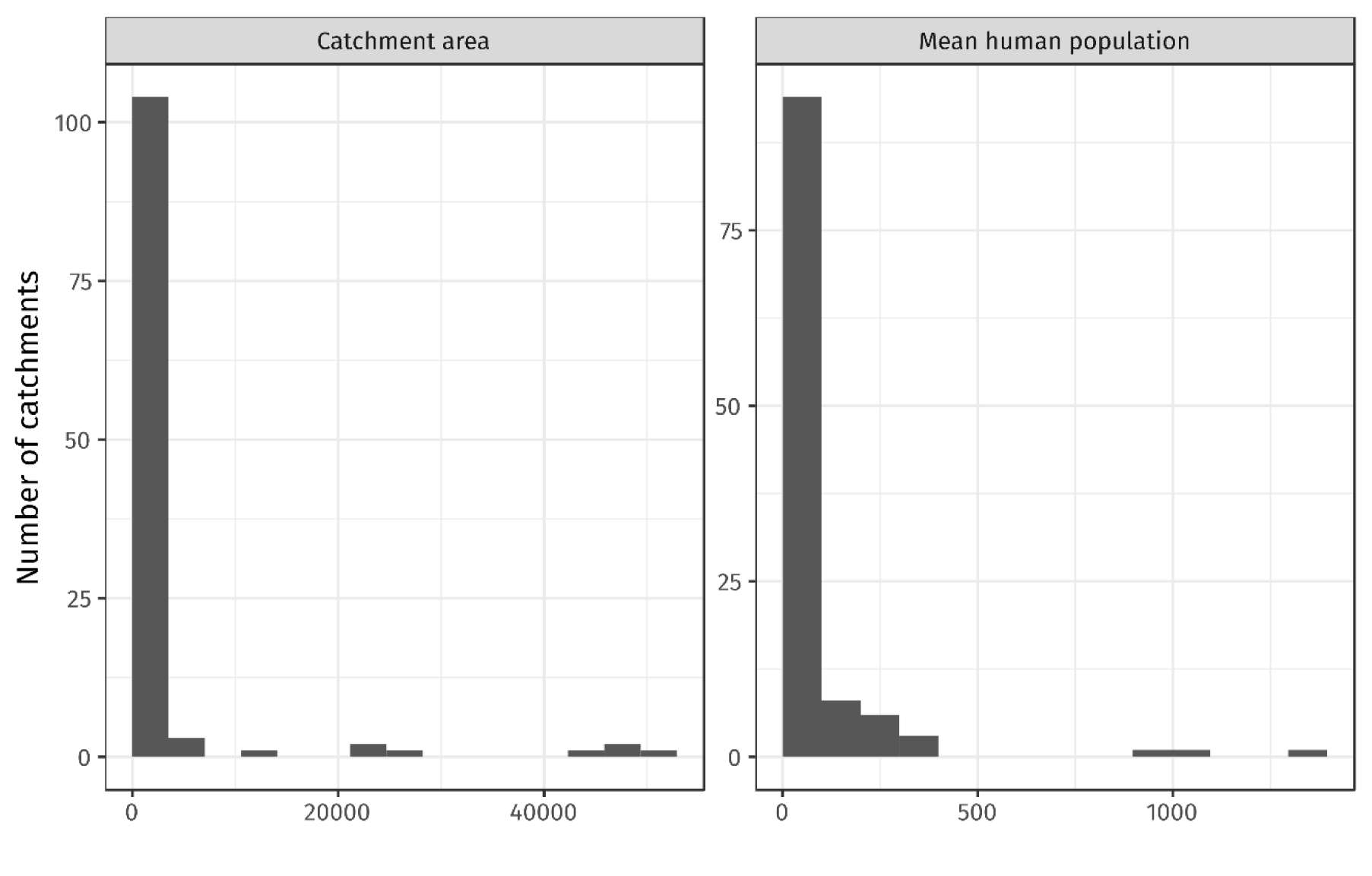
Histogram of catchment area and mean human population size in the catchment.

### Predator’s reliance on aquatic subsidies

The contribution of aquatic subsidies was significantly higher than 50% (effect size = 0.07, CI 95%: 0.013 – 0.13: fig. 4). Our model accounted for 95.3% of the heterogeneity in diet (R^2^=95.3, Q= 207.5, df = 19, p < 0.001) with a significant overall effect of moderators (Q_M_ =272.7, df = 23, p < 0.001). Sampling year and freshwater ecoregion both had a significant effect (Q_M_ = 76.4, df = 4, p < 0.001 for year and Q_M_ = 168.8, df = 15, p < 0.001 for ecoregion). The predator taxonomic group effect was not significant (Q_M_ = 7.88, df = 4, p = 0.096), whereas the test for residual heterogeneity was significant (Q_E_ = 63.5, df = 3, p < 0.0001), and most of the unaccounted variance is due to residual heterogeneity (I2= 95.3%).

**Figure 4:**
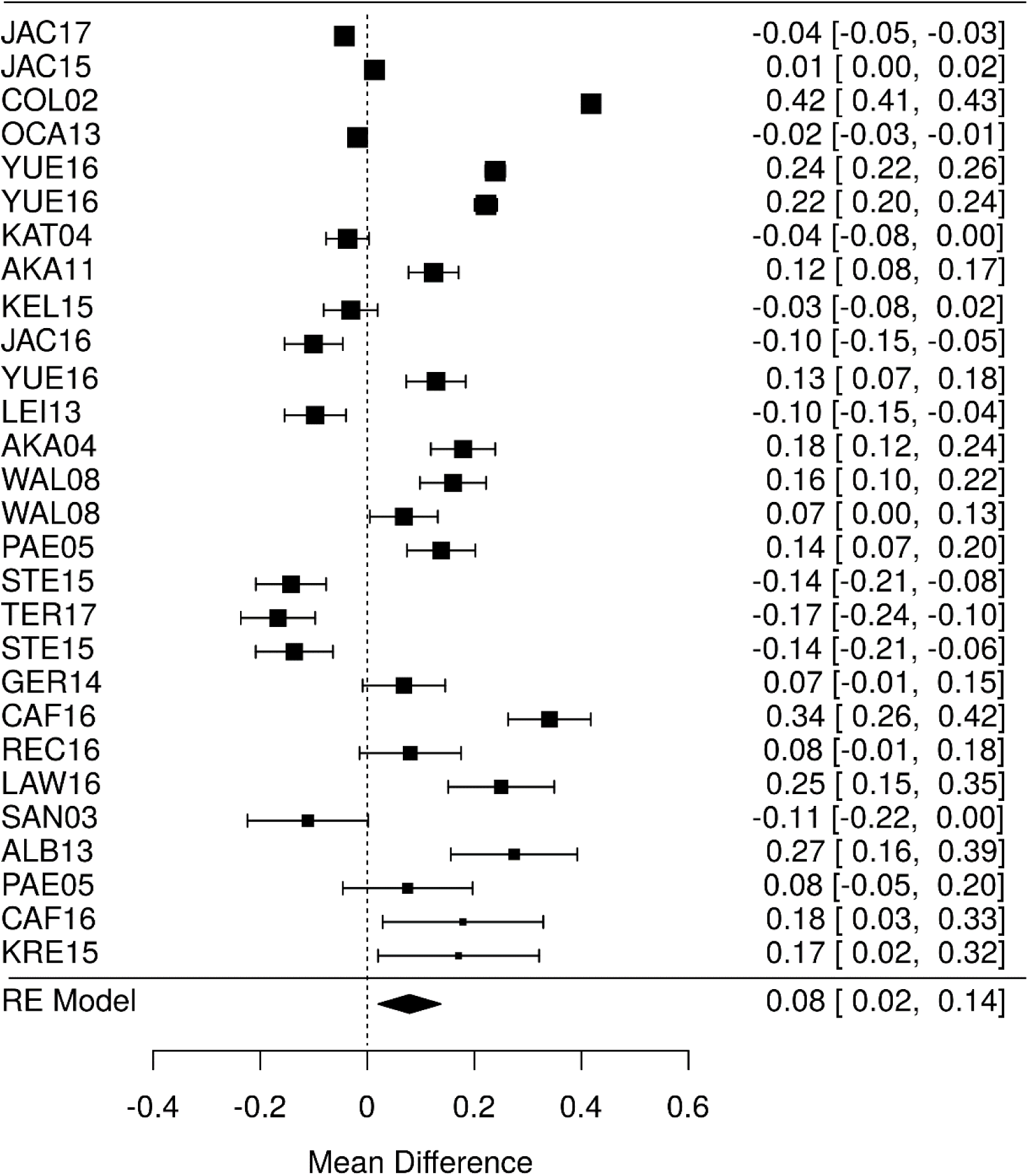
Forest plot showing the overall effect-size (observed proportion of aquatic prey in diet minus 0.5). Squares and bars denote means and 95% confidence intervals of the effect sizes, while the size of the squares reflects the weight of each study. Single studies are coded according to Table 1.

### Predictors of aquatic subsidies contribution

In the PLS regression model for spiders (two components: R^2^ = 0.394 and R^2^ = 0.460), the mean human population at both local scale and landscape scale as well as the percentage of agriculture at the landscape scale were the most important variables related to a high proportion of aquatic prey. In contrast, the percentage of non-forested natural habitats and open waters were related to low percentage of aquatic prey (fig 5). Despite high loading value, the percentage of open waters was weakly correlated to the percent of aquatic prey in the diet.

In the PLS regression model for carabid beetles (two components: R^2^ = 0.112 and R^2^ = 0.041), percent tree cover, forests, and water bodies at the landscape scale were the most important variables for low proportion of aquatic prey. The percentage of non-forested natural habitats, urban areas and agriculture at the landscape scale and the river width of the local scale were most important variables for high proportion of aquatic prey (fig. 6).

**Figure 5:**
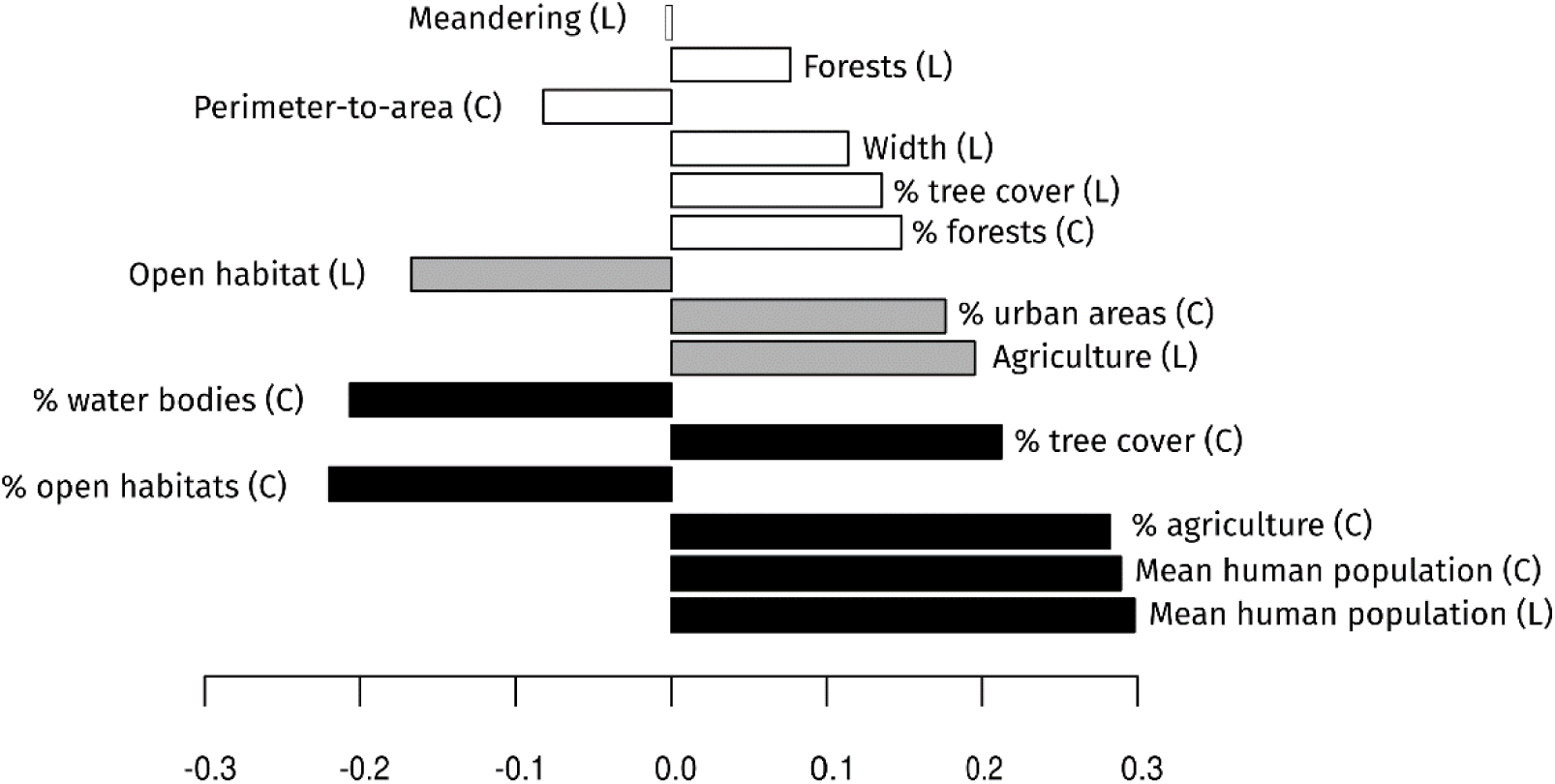
The variable weights of the first component in the PLS models for proportion of aquatic prey in spider diet. Positive weights indicate a positive relationship between the predictor and response variables and vice versa. Variables white bars are non-significant (VIP < 0.7). Variables with grey bars are significant with low explicative power (0.8 < VIP < 1). Variables in black are significant and are the most contributing variables (VIP > 1).

**Figure 6:**
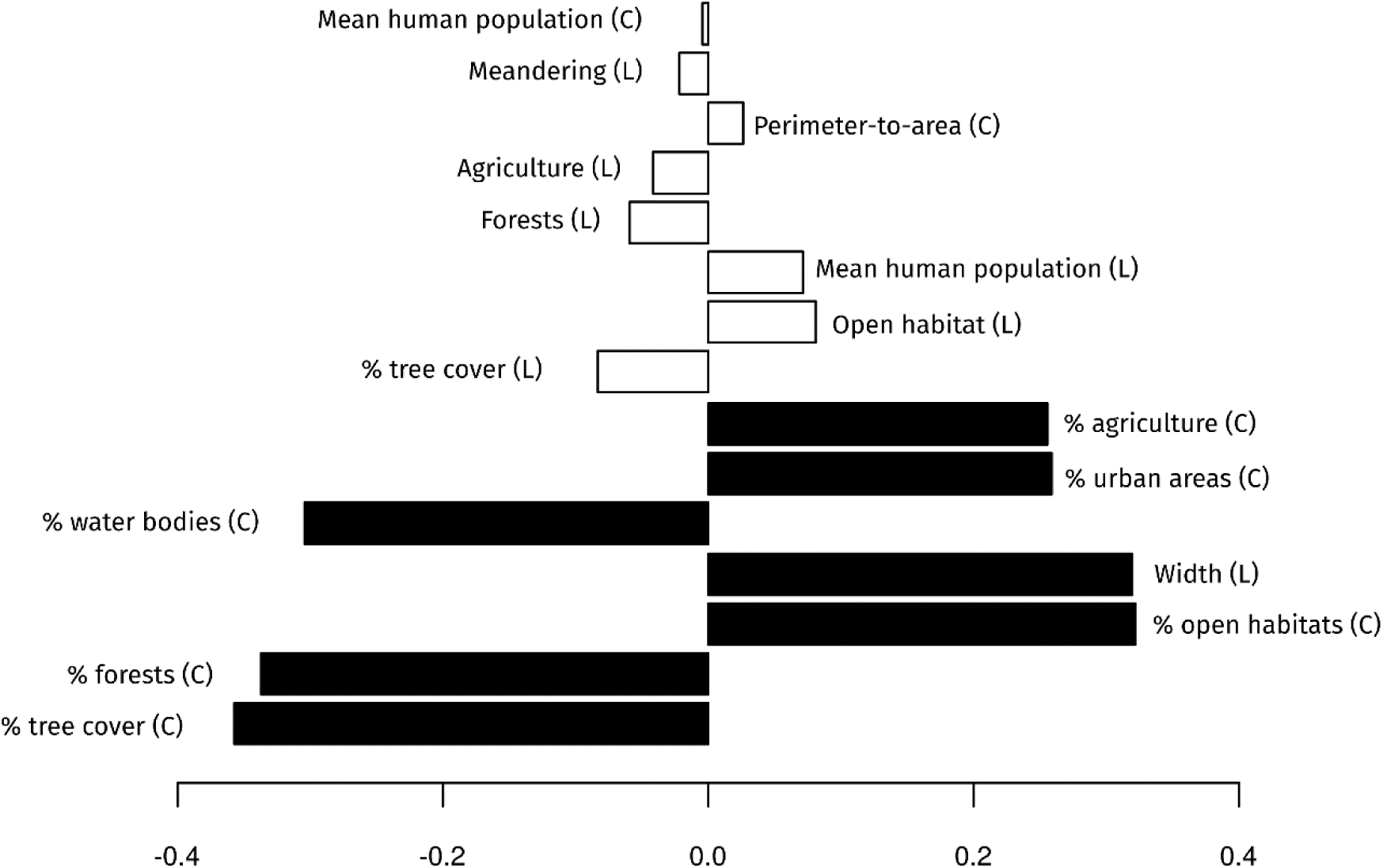
The variable weights of the first component in the PLS models for proportion of aquatic prey in the carabid beetle diet. Positive weights indicate a positive relationship between the predictor and response variables and vice versa. Variables white bars are non-significant (VIP < 0.7). Variables with grey bars are significant with low explicative power (0.8 < VIP < 1). Variables in black are significant and are the most contributing variables (VIP > 1).

## Discussion

Our study extends recent findings that demonstrate high levels of aquatic-to-terrestrial subsidies in riparian ecosystems (Bartels et al. 2012), improving both the resolution of subsidy quantification, and allowing inferences at broader ecological scales. Our meta-analysis also provides some of the strongest evidence to date of widespread effects of anthropogenic land use on the riparian food webs. These effects seem to be prevalent at the landscape scale, probably the most relevant scale for understanding the role of aquatic-terrestrial linkages for land management practices, such as proposed land use conversion or biodiversity conservation (Carpenter and Biggs 2009). Despite the general pattern of high aquatic subsidies use by terrestrial predators, we also documented significant inter-annual and geographic variations in these subsidies, largely driven by hydrologic cycles and ecoregion, respectively.

We found the diet of riparian predators to be highly dependent on aquatic subsidies (> 50%, overall effect size = 0.07). Since we re-computed diet partitioning from raw data to reduce mixing-model and discrimination-factor biases (Bond and Diamond 2011), our estimate is likely the most robust to date. This suggests that, in general, the proportion of aquatic subsidies in predator diets may be even higher than the 40% reported in Bartels et al.’s (2012) meta-analysis. We could not find any significant effect of predator taxonomic group, which might be due to the small number of studies dealing with groups other than carabid beetles and spiders. Given the wide geographic spread of our analysis and the pattern of high proportion of aquatic-derived carbon across the study sites, it seems likely that most predator taxa in riparian systems rely on these subsidies for more than 50% of their diet.

Perhaps unsurprisingly, we also found significant temporal (inter-annual) and spatial (ecoregion) variation in aquatic-to-terrestrial subsidies across the broad geographic scale of our study. Inter-annual climate-driven effects on stream hydrology (droughts vs floods) may have important impacts on aquatic and riparian communities (Power et al. 2008, Lafage et al. 2015b, Lafage and Pétillon 2016), and on aquatic and terrestrial food webs (Marks et al. 2000, O’Callaghan et al. 2013). Thus, inter-annual variation in hydrologic conditions act as a filter on functional traits of species and determines e.g. functional length of the riparian food chains. The significant effect of ecoregion on aquatic-to-terrestrial subsidies is probably due to region-specific differences in species communities, driven by both physical and ecological processes (Abell et al. 2008). It has been suggested that aquatic subsidy composition (especially through changes in species traits) is a key factor for resource use in the recipient system (Stenroth et al., 2015). Also, changes in predator communities might result in changes in species richness and functional diversity affecting the ability of predators to capture aquatic preys (e.g. for birds: Philpott et al. 2009).

Numerous studies have demonstrated the importance of landscape-scale processes on ecological status (e.g. Allan, 2004) and macro-invertebrate communities (aquatic: Lammert and Allan, 1999; Richards et al., 1996, terrestrial: Hendrickx et al., 2007; Lafage et al., 2015a). The relative importance of landscape- versus local-scale factors, however, is still under debate (Sandin and K. Johnson 2004, Stoll et al. 2016). In our study, the proportion of aquatic subsidies in terrestrial predator diets was almost exclusively related to landscape scale variables; the only significant local variable being human population. This was surprising, as many studies have highlighted the role of local vegetation (Tagwireyi and Sullivan, 2016), land use (Stenroth et al., 2015) and stream morphology (Iwata 2007, Muehlbauer et al. 2014). Our results could be related to the low resolution of our vegetation-related local variables, which were extracted from satellite data within a 100 m buffer. Nevertheless, variables related to stream morphology were not selected, although habitat geometry has been found to be the best predictor of trophic flow rate across habitat boundaries (Polis et al. 1997a).

At the landscape scale, ecosystem size did not explain the proportion of aquatic-terrestrial subsidies in predator diets. This may be due to the fact that the importance of ecosystem size and the direction of its relationship to predator diets can be system-specific, as conflicting relationships have been reported (Iwata, 2007, Stenroth et al. 2015, Jackson and Sullivan 2017). In our study, agricultural land use and urbanization, however, did have strong and consistent effects on terrestrial consumer diet which might be driven by either direct or indirect effects. First, by decreasing water quality, agriculture and urbanization usually directly affect the composition and quantity of aquatic subsidies (Carlson et al. 2016), shifting towards more and smaller species and resulting in better prey availability for smaller terrestrial predators (Stenroth et al. 2015). Second, land use changes may affect the amount and quality of terrestrial-to aquatic subsidies thereby indirectly influencing reciprocal aquatic-to-terrestrial subsidies (Nakano et al. 1999, Krell et al. 2015).

Habitat openness had opposite effects on spider and carabid diets so that spiders relied more on aquatic subsidies in forested catchments whereas carabids did the opposite. Riparian carabid beetles are usually small flattened winged species (O’Callaghan et al. 2013) more likely to capture small preys favored by open habitat (Carlson et al. 2016). Conversely, typical riparian spiders in forested catchment are large web-building spiders (e.g. *Tetragnatha* sp.) that are able to catch and consume large flying preys favored by forested habitats. Several studies have highlighted body size-trophic level linkages (e.g. (Cohen et al. 2003). A positive relationship between prey body-size and *Tetragnatha* use of aquatic subsidies has been previously demonstrated (Tagwireyi and Sullivan 2015).

Finally, both groups’ uses of aquatic subsidies were negatively related to the percentage of lakes at the landscape scale. Jonsson et al. (2018) recently found black fly larvae autochthony to be positively related to the lake proportion in river. In our case, it is most likely a geographical artefact. Sites located in Sweden presented the largest proportion of lakes and the smallest proportion of aquatic subsidies in predator’s diet.

The literature on insect emergence is heavily biased towards small streams (Muehlbauer et al. 2014, Schindler and Smits 2017). We found the same pattern, plus a geographical bias, for studies on predators’ diet using stable isotopes. Most of the studies we used were located in the northern hemisphere, in small-forested catchments with low proportions of agriculture or urbanization (except for studies specifically dealing with the impact of these land use related variables). As agriculture represents the main land use type in many developed catchments (Allan 2004) and urban land use exerts a disproportionately large influence on aquatic systems (Paul and Meyer 2001) we call for the development of studies on large rivers, and on catchment impacted by agriculture and urbanization. Studies are also needed on southern hemisphere streams.

Our study is the first worldwide meta-analyses to use exclusively stable isotope studies in order to better integrate the temporal component of terrestrial predator diets. We demonstrated a high reliance (more than 50%) of terrestrial predators on aquatic subsidies across broad geographic regions, despite large geographic and inter-annual variations.

We further demonstrated a large effect of anthropogenic land use at the catchment scale across geographic regions. Linking these two key findings suggests that more attention to broad-scale landscape patterns is warranted to improve our understanding of how these cross-boundary energy flows affects biodiversity and ecosystem function of tightly coupled aquatic-terrestrial systems.

## Acknowledgments

We would like to thank Karlstad University for funding this study within the framework of its strong research groups program. We particularly want to thank authors that sent us raw data: B.K. Jackson and K. Stenroth and Andrew C. Parnell for his help with Bayesian statistics.

